# Inhibition of *Glyoxalase 1* reduces alcohol self-administration in dependent and nondependent rats

**DOI:** 10.1101/230995

**Authors:** Giordano de Guglielmo, Dana E. Conlisk, Amanda M. Barkley-Levenson, Abraham A. Palmer, Olivier George

**Author notes:** Corresponding authors: Dr. Giordano de Guglielmo Department of Neuroscience, The Scripps Research Institute, 10550 North Torrey Pines Road, SP30-2400, La Jolla, CA 92037, USA., Dr. Olivier George Department of Neuroscience, The Scripps Research Institute, 10550 North Torrey Pines Road, SP30-2400, La Jolla, CA 92037, USA.

## Abstract

Previous studies showed that the glyoxalase 1 (*Glo1*) gene modulates anxiety-like behavior, seizure susceptibility, depression-like behavior, and alcohol drinking in the drinking-in-the-dark paradigm in nondependent mice. Administration of the small-molecule GLO1 inhibitor *S*-bromobenzylglutathione cyclopentyl diester (pBBG) decreased alcohol drinking in nondependent mice, suggesting a possible therapeutic strategy. However, the preclinical therapeutic efficacy of pBBG in animal models of alcohol dependence remains to be demonstrated. We tested the effect of pBBG (7.5 and 25 mg/kg) on operant alcohol self-administration in alcohol-dependent and nondependent rats. Wistar rats were trained to self-administer 10% alcohol (v/v) and made dependent by chronic intermittent passive exposure to alcohol vapor for 5 weeks. Pretreatment with pBBG dose-dependently reduced alcohol self-administration in both nondependent and dependent animals, without affecting water self-administration. pBBG treatment was more effective in dependent rats than in nondependent rats. These data extend previous findings that implicated *Glo1* in alcohol drinking in nondependent mice by showing even more profound effects in alcohol-dependent rats. These results suggest that the pharmacological inhibition of GLO1 is a relevant therapeutic target for the treatment of alcohol use disorders.

**Highlights:**

- Alcohol use disorder (AUD) places an enormous burden on society, and there is an urgent need for new druggable targets.
- Glo1 inhibition by pBBG dose-dependently reduces alcohol self-administration in both nondependent and dependent animals.
- pBBG treatment is more effective in reducing alcohol intake in dependent rats than in nondependent rats.

## 1 Introduction

Alcohol use disorder (AUD) is a chronic relapsing disorder that is characterized by compulsive alcohol use (Koob and Le Moal, 1997), which places an enormous burden on society. In addition to its psychological and societal toll, AUD was estimated to cost the United States’ economy USD$249 billion in 2010 alone (Sacks et al., 2015). Current behavioral and pharmacological treatments have limited efficacy, and there is an urgent need for new and more effective treatments (Koob, 2010). Alcohol use disorder has high comorbidity with several psychiatric disorders, including generalized anxiety disorder and depression. Thus, the development of treatments for AUD that may also address psychiatric comorbidities is critically important.

Voluntary binge-like alcohol drinking was recently shown to be modulated by glyoxalase 1 (*Glo1*) gene expression (McMurray et al., 2017b). The overexpression of *Glo1* increased drinking, and genetic knockdown of *Glo1* and pharmacological inhibition of the enzyme (GLO1) that is encoded by *Glo1* decreased alcohol drinking in nondependent mice in the drinking-in-the-dark (DID) paradigm (McMurray et al., 2017b). Interestingly, a role for *Glo1* in two other behavioral domains that are often comorbid with alcoholism has also been demonstrated. The genetic and pharmacological manipulation of *Glo1* also affects anxiety-like behavior (Distler et al., 2012; McMurray et al., 2016; Williams et al., 2009) and depression-related behavior (McMurray et al., 2017b). This raises the possibility that GLO1 inhibitors may be useful for the treatment of AUD and other common comorbid conditions. However, the effect of small-molecule inhibitors of GLO1 on alcohol self-administration in animal models of alcohol dependence remains to be demonstrated.

The present study tested the effect of the small-molecule GLO1 inhibitor *S*-bromobenzylglutathione cyclopentyl diester (pBBG) on alcohol self-administration in alcohol-dependent and nondependent rats. We used the chronic intermittent ethanol (CIE) model combined with operant self-administration, a model that has been shown to have robust predictive validity for alcoholism and construct validity for the neurobiological mechanisms of alcohol dependence (Heilig and Koob, 2007; Koob, 2009). Rats that are made dependent by CIE exhibit clinically relevant blood alcohol levels (BALs; 150-250 mg/100 ml), an increase in alcohol drinking when tested during early and protracted abstinence, and compulsive-like alcohol drinking (e.g., responding despite adverse consequences; Kimbrough et al., 2017b; Leao et al., 2015; O’Dell et al., 2004; Roberts et al., 1996; Schulteis et al., 1995; Vendruscolo et al., 2012). Alcohol dependence that is induced by alcohol vapor results in withdrawal symptoms during both acute withdrawal and protracted abstinence (de Guglielmo et al., 2017; Kallupi et al., 2014; Macey et al., 1996; Vendruscolo and Roberts, 2014), anxiety-like behavior (Valdez et al., 2002), irritability-like behavior (Kimbrough et al., 2017a), and the development of mechanical hyperalgesia (Edwards et al., 2012). We hypothesized that if GLO1 is a relevant target for the treatment of AUD, then pBBG treatment will reduce alcohol self-administration in both dependent and nondependent rats.

## 2 Materials and Methods

### *2.l* Animals

Adult male Wistar rats (*n* = 12; Charles River, Raleigh, NC, USA), weighing 225-275 g at the beginning of the experiments, were housed in groups of two per cage in a temperature-controlled (22°C) vivarium on a 12 h/12 h light/dark cycle (lights on at 8:00 PM) with *ad libitum* access to food and water. All of the behavioral tests were conducted during the dark phase of the light/dark cycle. All of the procedures adhered to the National Institutes of Health Guide for the Care and Use of Laboratory Animals and were approved by the Institutional Animal Care and Use Committee of The Scripps Research Institute.

### *2.2* Operant self-administration

Self-administration sessions were conducted in standard operant conditioning chambers (Med Associates, St. Albans, VT, USA). The animals were first trained to self-administer 10% alcohol (v/v) and water solutions until stable responding was maintained. First, to facilitate the acquisition of operant self-administration, the rats were initially provided free-choice access to 10% alcohol (v/v) and water for 1 day in their home cages to habituate them to the taste of alcohol. Second, the rats were subjected to an overnight session in the operant chambers with access to one lever (right lever) that delivered water on a fixed-ratio 1 (FR1) schedule of reinforcement. Food was available *ad libitum* during this training. Third, after 1 day off, the rats were subjected to a 2 h session on an FR1 schedule for 1 day and a 1 h session on an FR1 schedule the next day, with one lever delivering alcohol (right lever). All of the subsequent sessions lasted 30 min, and two levers were available (left lever: water; right lever: alcohol) until stable levels of intake were reached.

### *2.3* Alcohol vapor chambers

The rats were made dependent by chronic intermittent exposure to alcohol vapors as previously described (Gilpin et al., 2008; O’Dell et al., 2004). They underwent cycles of 14 h ON (BALs during vapor exposure ranged between 150 and 250 mg%) and 10 h OFF, during which behavioral testing for acute withdrawal occurred (i.e., 6-8 h after the vapor was turned OFF when brain and blood alcohol levels are negligible; Gilpin et al., 2009). In this model, rats exhibit somatic withdrawal signs and negative emotional symptoms, reflected by anxiety-like responses and elevated brain reward thresholds (de Guglielmo et al., 2016; Edwards et al., 2012; O’Dell et al., 2004; Rimondini et al., 2002; Schulteis et al., 1995).

### *2.4* Operant self-administration during alcohol vapor exposure

Behavioral testing during alcohol vapor exposure occurred three times per week. The rats were tested for alcohol (and water) self-administration on an FR1 schedule in 30-min sessions. Operant self-administration on an FR1 schedule requires minimal effort by the animal to obtain the reinforcement and herein was considered a measure of intake.

### *2.5* Drugs

The 10% alcohol (v/v) drinking solution was prepared by diluting 95% alcohol (v/v) in water. pBBG was synthesized in the laboratory of Prof. Alexander Arnold (University of Wisconsin, Milwaukee; McMurray et al., 2017a; McMurray et al., 2017b). pBBG was dissolved in vehicle (8% dimethylsulfoxide, 18% Tween-80, and 74% distilled water) and administered intraperitoneally 30 min before the test session.

### *2.6* Effect of systemic pBBG on alcohol self-administration in nondependent and dependent rats

Wistar rats (*n* = 12) were trained to self-administer 10% alcohol (v/v) and water until stable self-administration was established (± 10% over the last three sessions). The rats were then intraperitoneally injected with pBBG (0, 7.5, and 25 mg/kg/ml) according to a Latin-square design with a baseline alcohol self-administration session between tests. At the end of the treatment, the rats were re-baselined for alcohol self-administration to exclude possible long-lasting effects of the compound. After three self-administration tests, the animals were moved to the vapor chambers for 3 weeks. Blood samples were collected once weekly to determine BALs. The rats were then tested for alcohol (and water) self-administration on an FR1 schedule of reinforcement in 30-min sessions until the escalation of self-administration was observed. At this point, the treatment began, and the animals were intraperitoneally injected with pBBG (0, 7.5, and 25 mg/kg/ml) according to a Latin-square design with a baseline alcohol self-administration session between tests.

### *2.7* Statistical analysis

The data were analyzed using one-way repeated-measures analysis of variance (ANOVA), followed by the Newman Keuls *post hoc* test. The escalation data and changes *vs*. baseline were analyzed using Student’s *t*-test. Values of *p* < 0.05 were considered statistically significant.

## 3 Results

### *3.1* Systemic pBBG reduces alcohol self-administration in nondependent and dependent rats

pBBG significantly reduced alcohol self-administration in nondependent rats. This effect was confirmed by the one-way ANOVA, which revealed a significant effect of treatment (*F*_2,11_ = 9.53, *p* < 0.01). The Newman Keuls *post hoc* test showed that pBBG significantly reduced alcohol self-administration at the dose of 25 mg/kg (*p* < 0.01; Fig. 1A, a). The effect was dose-dependent, in which a significant difference was detected between the 7.5 and 25 mg/kg doses (*p* < 0.05; Fig. 1A, a). Water self-administration was unaffected by the treatment (*F*_2,11_ = 0.36, *p* > 0.05). Further analysis of the percent reduction of alcohol self-administration relative to baseline showed that 7.5 mg/kg pBBG did not affect alcohol intake (12.7% ± 10.4%), whereas 25 mg/kg significantly decreased alcohol self-administration (44.6% ± 13.4%; *p* < 0.01; Fig. 1B).

**Figure 1.**
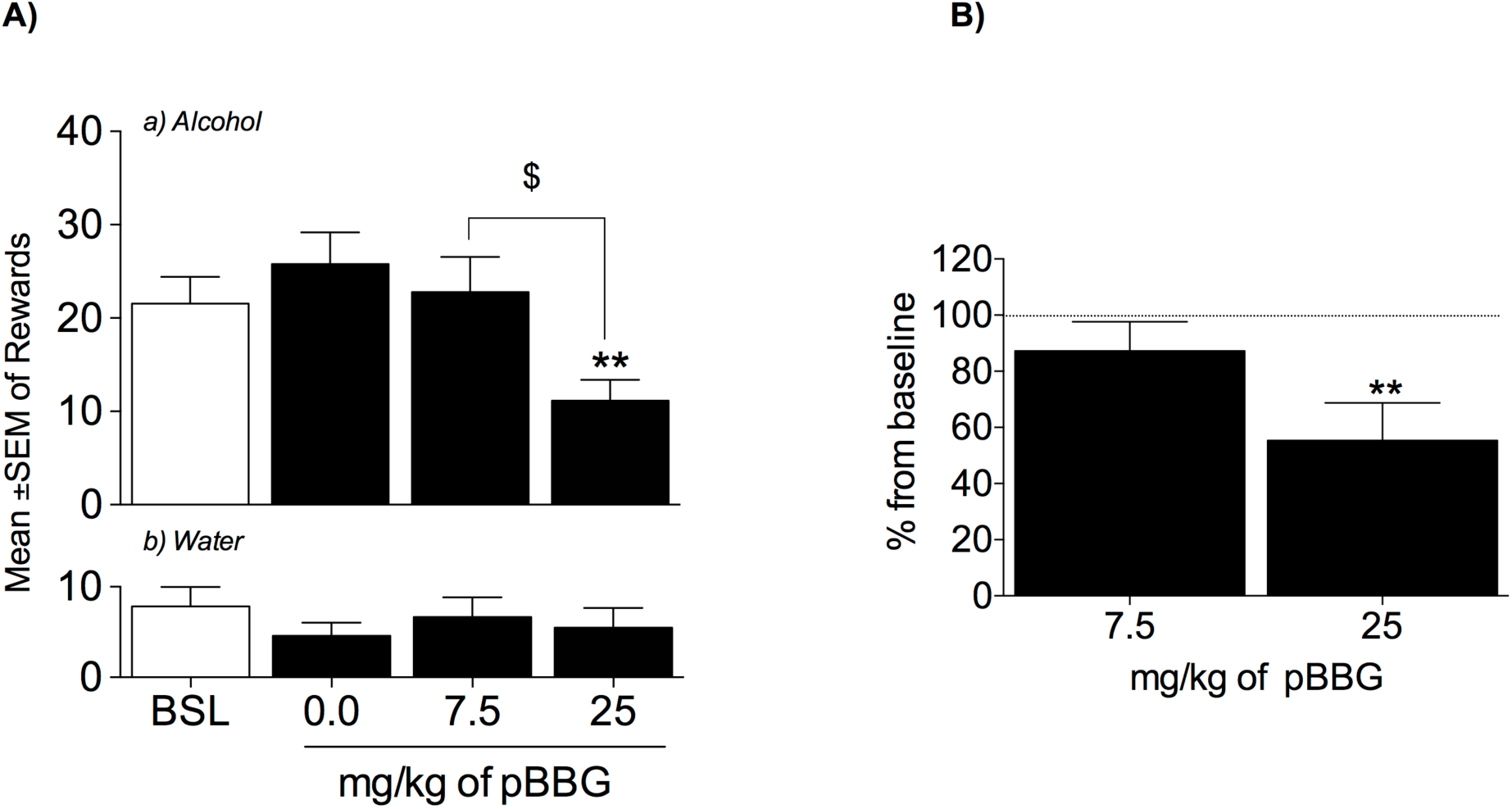
pBBG reduced alcohol self-administration in nondependent rats. (**A**) Effect of pBBG on (a) alcohol intake and (b) water intake. \*\**p* < 0.01, *vs*. 0.0 mg/kg; ^$^*p* < 0.05, *vs*. 7.5 mg/kg. (**B**) Percent reduction *vs*. baseline. \*\**p* < 0.01, vs. baseline.

After the re-stabilization of alcohol self-administration and before vapor exposure, the mean ± SEM number of reinforced responses for alcohol was 23.1 ± 2.6. After 6 weeks of alcohol exposure, the animals significantly escalated their alcohol intake, and the mean ± SEM number of reinforced responses increased to 49.3 ± 7.5. The one-way ANOVA revealed a significant effect of vapor exposure (*F*_8,11_ = 3.1, *p* < 0.05). The Newman Keuls *post hoc* test showed that the animals significantly increased their alcohol consumption after the sixth test session (*p* < 0.05; Fig. 2A). Blood alcohol levels significantly increased from week 1 to week 6 of vapor exposure (*t*-test: *t* = 10.10, df = 11, *p* < 0.001; Fig. 2B).

**Figure 2.**
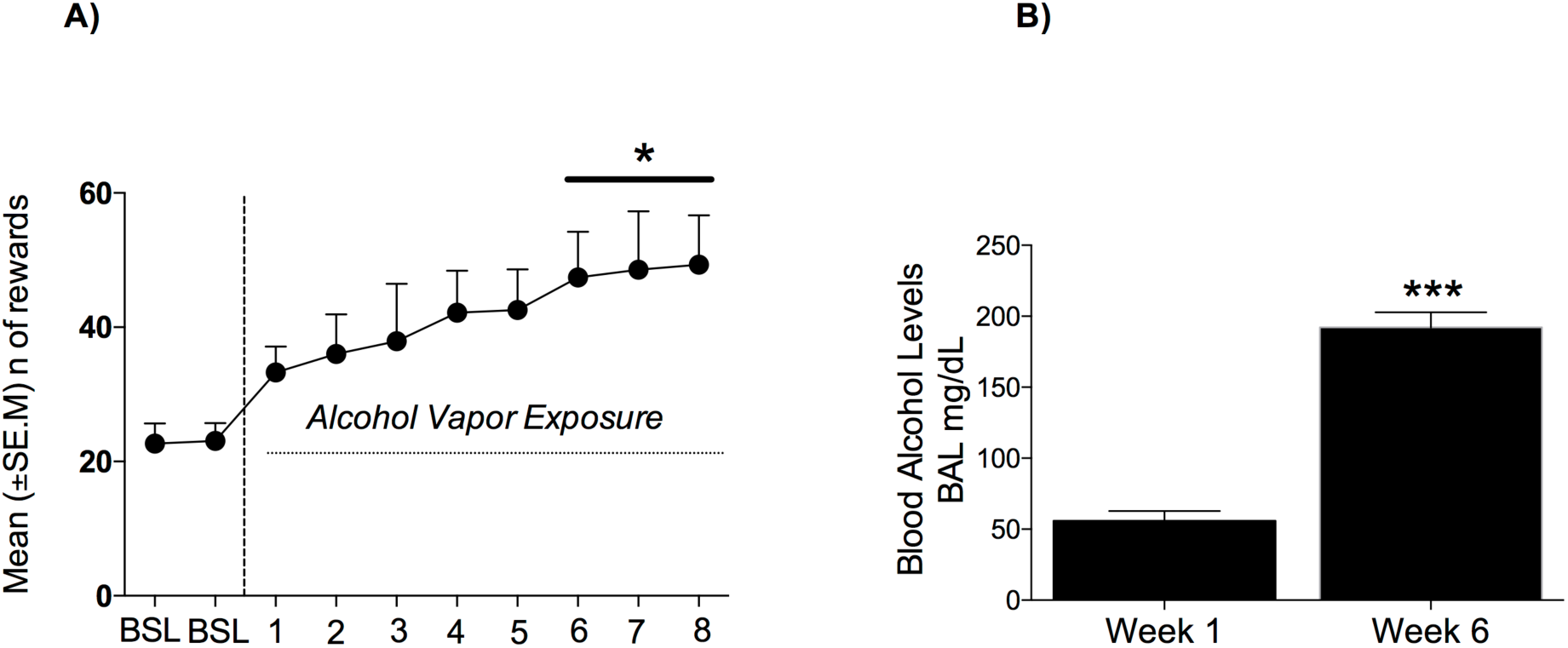
(**A**) Escalation of alcohol self-administration during passive alcohol vapor exposure. \**p* < 0.05, *vs*. last baseline day. (**B**) Blood alcohol levels after 1 and 6 weeks of passive alcohol vapor exposure. \*\*\**p* < 0.001, *vs*. week 1.

The intraperitoneal injection of pBBG significantly reduced alcohol self-administration in dependent rats (*F*_2,11_ = 16.72, *p* < 0.001). The Newman Keuls *post hoc* test showed that pBBG significantly reduced alcohol self-administration at both 7.5 mg/kg (*p* < 0.05) and 25 mg/kg (*p* < 0.01; Fig. 3A, a). The effect was dose-dependent, in which a significant difference was detected between the 7.5 and 25 mg/kg doses (*p* < 0.05; Fig. 3A, a). Water intake was unaffected by the treatment (*F*_2,11_ = 3.1, *p* > 0.05; Fig. 3A, b). Further analysis of the percent reduction of alcohol self-administration relative to alcohol intake before treatment showed that pBBG significantly decreased alcohol self-administration at both 7.5 mg/kg (40.7% ± 8.6%, *p* < 0.01) and 25 mg/kg (74.1% ± 9.3%, *p* < 0.001; Fig. 3B).

**Figure 3.**
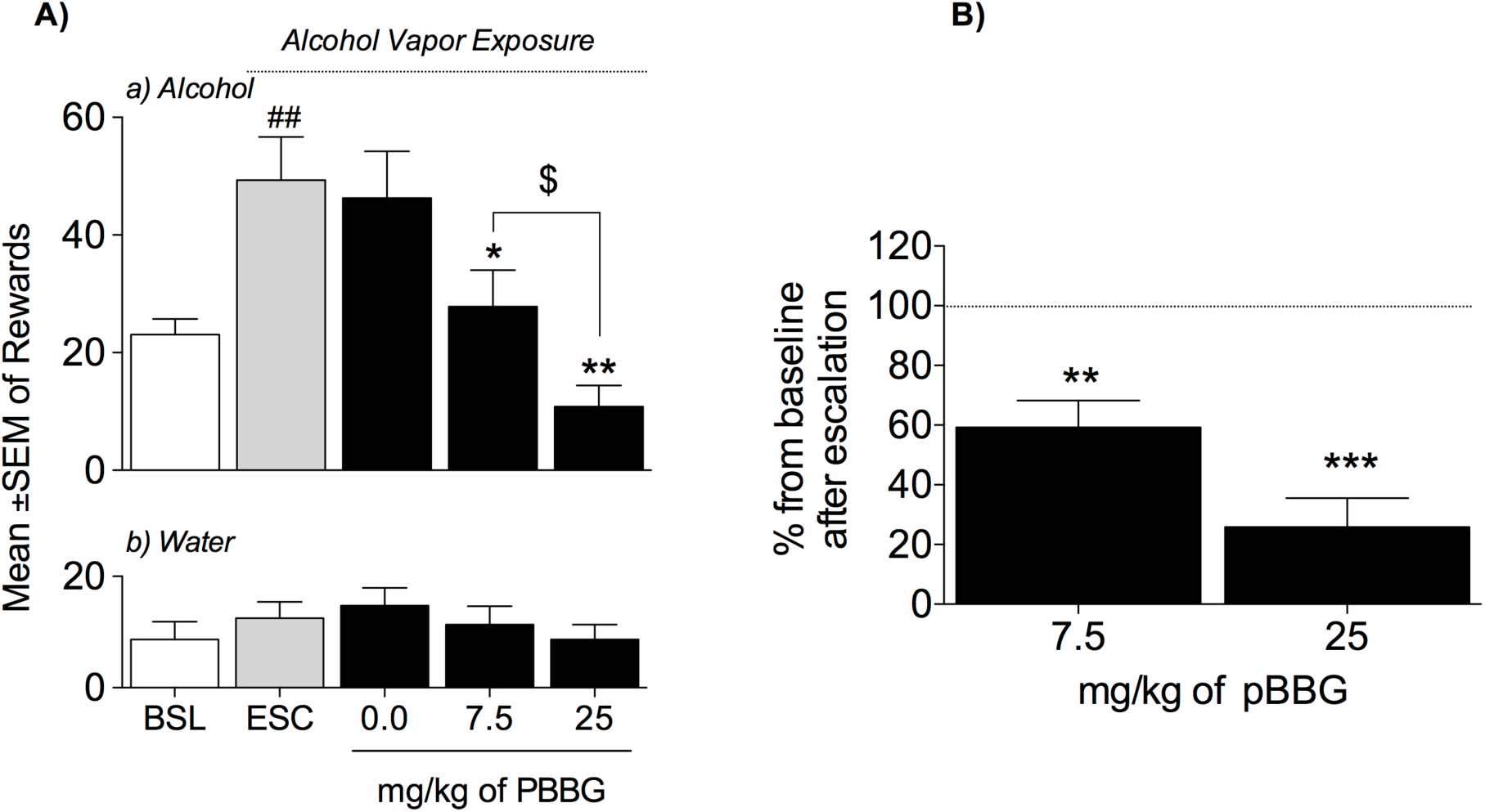
pBBG reduced alcohol self-administration in dependent rats. (**A**) Effect of pBBG on (a) *M-LL* alcohol intake and (b) water intake.^##^*p* < 0.01, *vs*. baseline; \**p* < 0.05, \*\**p* < 0.01, *vs*. 0.0 mg/kg; ^$^*p* < 0.05, *vs*. 7.5 mg/kg. (**B**) Percent reduction vs. baseline. \*\**p* < 0.01, \*\*\**p* < 0.001, *vs*. baseline.

## 4 Discussion

The present study demonstrated a role for GLO1 in the regulation of alcohol consumption in dependent rats and extended previous findings by showing a key role for GLO1 in modulating alcohol drinking in nondependent animals. A key novel finding was that pBBG, a small-molecule inhibitor of GLO1, reduced alcohol self-administration in dependent rats and was more effective in dependent rats than in nondependent rats. Treatment with pBBG (7.5 and 25 mg/kg) significantly reduced alcohol self-administration to baseline (predependent) levels. Water consumption was unaffected by pBBG treatment, thus excluding possible nonspecific effects of pBBG on alertness and locomotor activity that may have biased the results. These results are consistent with previous studies that found that similar doses of pBBG had no effect on water, saccharin, or quinine drinking (McMurray et al., 2017b), did not exert a sedative effect as measured by the loss-of-righting reflex, and did not cause ataxia as measured by the balance beam task (McMurray et al., 2017b). Recent work has also shown that GLO1 manipulation does not alter the locomotor stimulant or depressant effects of alcohol, demonstrating that GLO1 inhibition does not reduce alcohol drinking simply by making alcohol more potent (Barkley-Levenson et al., unpublished results).

Electrophysiological studies showed that methylglyoxal (GLO1’s substrate) is a novel, endogenously produced, competitive partial agonist at γ-aminobutyric acid-A (GABA_A_) receptors (Distler et al., 2012). Therefore, changes in methylglyoxal concentrations that are caused by pBBG-induced GLO1 inhibition are presumed to modulate alcohol consumption through the actions of methylglyoxal at GABA_A_ receptors.

Alcohol does not have an identified specific neurotransmitter binding site, but ethanol receptive elements within the membrane may provide a sensitive site for ethanol actions (Tabakoff and Hoffman, 1992). Preclinical studies have identified several neuromodulatory systems that are modulated by ethanol and that represent potential targets for future medications (De Witte et al., 2005; Spanagel and Kiefer, 2008; Tanchuck et al., 2011; Funk et al., 2007; Gehlert et al., 2007;Valdez and Koob, 2004; Nealey et al., 2011; Walker et al., 2011; de Guglielmo et al., 2015; Martin-Fardon et al., 2010; Logrip et al., 2015; King et al., 2017; Gomez et al., 2015; Franklin et al., 2015; Hu et al., 2011; Logrip et al., 2014).

Many of the behavioral effects that are associated with alcohol use are mediated by the interactions between alcohol and the GABA system (Centanni et al., 2017; Lindemeyer et al., 2014; Roberto et al., 2003; Roberto et al., 2004; Weiner and Valenzuela, 2006), which has stimulated investigations of the effects of GABAergic drugs on alcohol self-administration in rats (Addolorato et al., 2012; Colombo et al., 2003; Stopponi et al., 2012) and humans (Addolorato et al., 2002; Brower et al., 2008; Mason et al., 2014). A recent study found that the GABA analogue gabapentin (Neurontin; Pfizer) dose-dependently reduced alcohol intake in postdependent rats (Roberto et al., 2008). In the same study, electrophysiological experiments showed that the application of gabapentin in central nucleus of the amygdala (CeA) slice preparations from postdependent rats reduced the alcohol-induced increase in GABA currents (Roberto et al., 2008). Conversely, gabapentin produced robust increases in inhibitory transmission in the CeA in nondependent animals, an effect that was accompanied by a nonsignificant trend toward an intra-CeA gabapentin-induced increase in alcohol intake in nondependent rats, which is consistent with the opposite electrophysiological profiles of gabapentin in alcohol-dependent and -naive rats (Roberto et al., 2008). These results support the hypothesis that a history of alcohol exposure may produce long-term neuroadaptations that ultimately lead to excessive alcohol consumption (Heilig and Koob, 2007) and are consistent with our data that demonstrated greater efficacy of pBBG in alcohol-dependent animals. This hypothesis is also consistent with previous studies that evaluated the GABA modulator Pregabalin (Lyrica; Pfizer). Pregabalin is a structural analogue of GABA that is approved by the United States Food and Drug Administration for the treatment of partial epilepsy, neuropathic pain, and more recently generalized anxiety disorder. Treatment with pregabalin selectively reduced home-cage alcohol drinking, alcohol self-administration, and the stress- and cue-induced reinstatement of alcohol seeking in Marchigian Sardinian alcohol-preferring (msP) rats (Stopponi et al., 2012), a strain that has several behavioral and neurobiological similarities to postdependent rats (Ciccocioppo et al., 2006; Gehlert et al., 2007; Hansson et al., 2006). Thus, there is ample evidence of a role for GABA in alcohol drinking, and our results further indicate that GABA modulators may be a valid strategy for the management of AUDs.

In summary, the present study demonstrated the preclinical therapeutic efficacy of pBBG in an animal model of escalated alcohol drinking in dependent rats and extended previous findings that implicated *Glo1* in alcohol drinking in nondependent mice. These results suggest that GLO1 is a relevant therapeutic target for the treatment of AUDs that should be further investigated.

## Authors Contributions

GdG, AAP, and OG were responsible for the study concept and design. GdG, DEC, and ABL contributed to the acquisition of animal data. GdG and OG drafted the manuscript. All authors critically reviewed the content and approved the final version for publication.

## Acknowledgements

The authors thank Michael Arends for proofreading the manuscript.

